# The metabolic resistance blueprint: Genomic dissection of DDT resistance in historical *kdr*-free East African *Anopheles gambiae*

**DOI:** 10.64898/2026.02.15.705978

**Authors:** Talal Al-Yazeedi, Marion Morris, Abdullahi Muhammad, Omer S. Alkhnbashi, Naomi A. Dyer, Hilary Ranson

**Affiliations:** Liverpool School of Tropical Medicine, Liverpool, UK; Khalifa Centre for Genetic Engineering and Biotechnology, United Arab Emirates University, P.O. Box 15551, Al Ain, UAE; College of Medicine, Mohammed Bin Rashid University of Medicine and Health Sciences (MBRU), Dubai Healthcare City, Dubai, United Arab Emirates

## Abstract

Insecticide resistance in *Anopheles gambiae* poses a significant threat to malaria control and eradication efforts across sub-Saharan Africa. While target-site resistance mechanisms are well-characterised, the evolutionary origins of metabolic resistance remain poorly understood. We employed bulk segregant analysis (BSA) using unique historical genetic crosses established in the late 1990s between a DDT-resistant ZAN/U strain (Originally from Zanzibar) and a susceptible strain to map the genomic architecture of early metabolic resistance. Critically, the ZAN/U strain exhibited DDT resistance without *kdr* mutations, providing an ideal genetic background for isolating glutathione S-transferase epsilon (*GSTe*)-mediated resistance mechanisms that evolved during the DDT control program era. BSA analysis revealed a major quantitative trait locus on chromosome 3R spanning the *GSTe* gene cluster (*GSTe*1-8). BSA identified 20 significant amino acid substitutions distributed across all eight *GSTe* genes, in addition to variations in the promoter region of *GSTe2* and *GSTe3*, which were previously determined to be overexpressed in the ZAN/U strain. East African field-collected samples from the MalariaGEN Ag1000G project confirmed that many of these mutations are under long-term selection pressure in natural populations. This work provides a characterisation of the *GSTe* cluster’s role in establishing the metabolic resistance foundation that continues to compromise vector control efforts today. Additionally, BSA analysis was performed on progeny from crosses established in early 2000s between a permethrin-resistant RSP-ST strain originating from Western Kenya and a susceptible strain, identifying expected target-site resistance at the *vgsc* locus.

## Introduction

Malaria remains one of the world’s deadliest infectious diseases, with an estimated 263 million cases and 597,000 deaths in 2023, with 94% of cases and 95% of deaths occurring in Sub-Saharan Africa ^1^. Children under 5 accounted for about 76% of all malaria deaths, with a child dying nearly every minute from this preventable, treatable disease ^1^. Vector control through insecticide-treated nets (ITNs) represents the primary defence against malaria transmission and has averted millions of deaths since the year 2000 ^2^, yet this strategy faces an escalating threat from insecticide resistance in *Anopheles gambiae*, the principal malaria vector in sub-Saharan Africa ^3,4^. Insecticide resistance in *An. gambiae* operates through two primary mechanisms: target-site resistance and metabolic resistance ^5^. Mutations in the voltage-gated sodium channel, the target of both dichlorodiphenyltrichloroethane (DDT) and pyrethroid insecticides, cause knockdown resistance (*kdr*) ^6–9^, and readily tracked through PCR. However, metabolic resistance has a greater operational impact on malaria control ^8,10^. Metabolic resistance involves changes in the rate of sequestration, detoxification and transport of insecticides with increased expression of detoxification genes or a change in the protein conformation of these genes due to amino acid changes, enhancing the detoxification efficiency ^11^.

The glutathione S-transferases (*GSTs*) are important insecticide detoxification enzymes, catalysing the conjugation of glutathione to hydrophobic insecticides and the dehydrochlorination of DDT, facilitating their elimination from mosquito tissues ^12–14^. Although metabolic resistance to pyrethroids is more commonly conferred by elevated cytochrome P450 activity, GSTs have been associated with pyrethroid resistance ^15,16^ and hence understanding the evolutionary dynamics and mutational landscape of the *GSTs* can assist in predicting and managing contemporary resistance patterns.

The evolution of insecticide resistance in African malaria vectors reflects decades of changing vector control strategies. Dichlorodiphenyltrichloroethane (DDT) was developed in the 1940s and initially used with great effect in the WHO’s malaria campaigns of the 1950s and 1960s ^17,18^. However, DDT resistance was widespread by the early 1970s, emerging after several years of intensive application ^18,19^.

Synthetic pyrethroids were introduced in the 1990s in Indoor Residual Spraying to control DDT-resistant mosquitoes, but the initial wave of pyrethroid resistance was primarily a re-selection of old DDT resistance mechanisms, as both insecticide classes share a common sodium channel target. DDT is now rarely used for malaria control in Africa due to concerns about its environmental persistence. Following successful trials in the late 1990s, the use of ITNs was dramatically scaled up in the 21^st^ century, with over 2 billion ITNs containing pyrethroid insecticides distributed across the African continent in the last 25 years ^2,20^. Today, due to continuous exposure to insecticides via malaria control efforts and agricultural use, there is almost no African country with a fully pyrethroid-susceptible malaria vector ^1^.

The association of GSTs with DDT resistance and cytochrome P450S with pyrethroid resistance were reported in the early 1990s ^21^; however, the comprehensive study of these enzyme families was not possible until the complete *An. gambiae* genome was sequenced in 2002 ^22^. The *An. gambiae* genome contains more than 110 P450 genes, with clusters of CYP6 genes most widely associated with pyrethroid resistance in *An gambiae* ^23^. Multiple classes of GST genes are organised into distinct clusters across the genome. The eight genes in the epsilon class (*GSTe1-8*) are clustered on chromosome 3R and are the most important GST family for insecticide detoxification in *Anopheles* ^13,24,25^. In this study, we revisited mosquito samples from *An. gambiae* genetic crosses that were preserved for more than 20 years before being reused in this study. These genetic crosses were used in two separate studies to map resistance loci to DDT and permethrin using microsatellite markers. Genetic crosses were established between the DDT-resistant ZAN/U strain colonised from a DDT-resistant *An. gambiae* field population from Zanzibar, Tanzania, in 1982 and an insecticide-susceptible FAST strain (developed by selectively breeding the 4Ar/r strain of *An. gambiae* by Zheng et al. (1997) to obtain mosquitoes with the standard chromosome arrangement ^25,26^. The DDT-resistant ZAN/U strain lacked the *kdr* mutation, which made it an excellent model to isolate the molecular mechanism mediating metabolic resistance. QTL mapping with microsatellite markers identified two quantitative trait loci (QTL), a major resistance locus *rtd1* on chromosome 3 and *rtd2* on chromosome 2L, with an additive genetic effect ^25^. *Rtd1* was subsequently linked to the overexpression of a *GSTe* cluster, and signatures of selection at this locus have been reported in many African regions, correlating with resistance ^10,27^.

A similar approach was used to identify loci associated with pyrethroid resistance from a genetic cross between the FAST5 susceptible strain and the RSP-ST (reduced susceptibility to permethrin) strain ^28^). The RSP-ST strain originated from western Kenya and was derived from females collected in 1992, during an early trial of permethrin-impregnated bednets and curtains ^29,30^. Genotyping the RSP-ST strain at the sodium channel locus confirmed that the strain was fixed for the (L1014F) *kdr* mutation ^9^, and this mutation explained the first of the two QTL loci identified in this cross. The second maps to a region of the genome containing a large cluster of cytochrome P450 genes implicated in pyrethroid resistance ^9,31^.

While microsatellite markers successfully enabled QTL mapping of resistance loci, they provide only low-resolution mapping and cannot detect polymorphisms in candidate regions. The advancement of sequencing technology enabled the complete resequencing of thousands of laboratory and field-collected *An. gambiae* specimens, resulting in the discovery of genomic regions under insecticide selection pressure while identifying polymorphism in these regions at the SNP resolution. In this study, we performed whole-genome sequencing on pools of DNA extracted from mosquito samples from genetic crosses preserved for more than 20 years. We performed bulk segregant analysis (BSA) to map QTLs associated with resistance and uncover polymorphisms within these QTLs.

## Results

### BSA provides a rapid and cost-efficient approach to detecting genomic loci associated with resistance

BSA was conducted for each pairing of alive and dead DNA pools constructed from the crossing families ^25,28^. Single pair crosses of the DDT-resistant ZAN/U strain and the FAST5 susceptible strain were reared until the F8 generation. At the F8, progeny were exposed to DDT to separate individuals into resistant (alive) and susceptible (dead) progeny. For each of the two separate original crosses (DDT-A and DDT-B), DNA pools were constructed from segregating progeny, with separate pools for resistant and susceptible individuals. Each pool consisted of DNA from 80 individuals (See methods for details) ^25^. To investigate the historical molecular markers driving permethrin resistance, we constructed DNA pools based on susceptibility to permethrin from the historical cross between the RSP-ST strain and the FAST5 susceptible strain ^28^. BSA investigated pools from two crossing families (Perm-A and Perm-B) phenotyped at the F4 and F5 generation offspring, respectively. Offspring from the original cross were exposed to permethrin for different durations to maximise phenotypic segregation between resistant and susceptible individuals. This arrangement generated four DNA pools, two alive (AL) and two dead (DE), per family, with varying sample sizes and exposure time (see method for details).

Genome-wide sequencing of these pooled samples successfully identified single-nucleotide polymorphisms (SNPs) across all families by aligning to the *An. gambiae* PEST 4 reference genome. BSA was conducted for each alive and dead pool pairing, yielding four comparisons per family for permethrin crosses and two for DDT crosses (Table S1). To ensure reproducibility and facilitate broader application, we developed a comprehensive pipeline from a collection of bash scripts https://github.com/alyazeeditalal/Poolseq_analysis that automates the entire BSA workflow from raw sequencing data to quantitative trait loci (QTL) identification and visualisation. This pipeline enables consistent analysis across different experimental designs and insecticide treatments (Figure 1 and Table S1). Sequencing of DNA pools yielded an average of 9.74×10^7^ reads per pool after removal of read duplicates and low quality reads, with an average mapping rate of 97.8%, a mean of 49.1x coverage using the *An. gambiae* PEST genome ^32^ and a mean mapping quality of 47.2. The pipeline identified a total of 8,137,996 SNPs after filtration of low-quality SNPs, retaining only high-quality, biallelic variants with a genotype quality (GQ) ≥ 99, and depth (DP) ≥ 100 (Table S1). The high-quality SNPs were utilised for the BSA to identify regions of interests associated with resistance in each crossing family.

**Figure 1.**
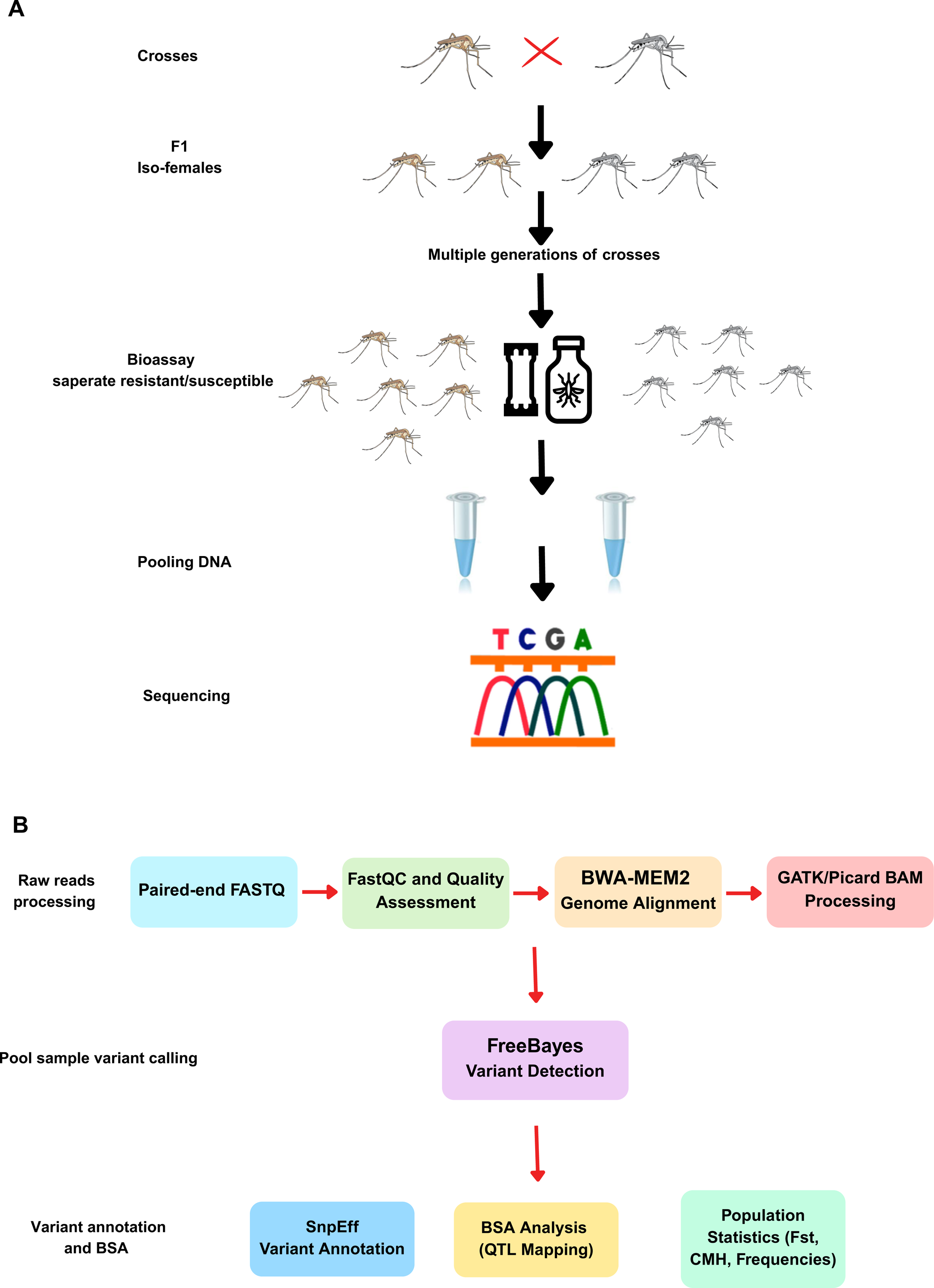
BSA analysis pipeline. **(A)** Bulk segregant analysis experimental design showing crossing scheme between resistant and susceptible strains. Single-pair crosses are established to produce F1 females. Impregnated F1 iso-females are separated, and their progeny are allowed to inter-cross through multiple generations to establish separate families. Progeny are exposed to insecticides and separated into resistant (alive) and susceptible (dead) phenotypic classes; DNA is then pooled within each class for sequencing. **(B)** Bioinformatics workflow for BSA analysis showing the complete pipeline from raw sequencing data to results. The pipeline encompasses paired-end sequencing quality control, genome alignment, BAM processing, variant calling, BSA analysis for QTL mapping, population statistics calculation, and variant annotation.

The significance level of the BSA QTL analysis was determined using several tests ^33^. First, the difference in SNP-index between alive and dead bulk (delta SNP-index) was calculated ^34^. Additionally, G-statistics (G value) were calculated genome-wide, and a smoothing method, G-prime (G’), was calculated on the G value in a sliding window of 3 Mb ^35^. To determine the significance of the detected QTL, p-values were calculated by comparing the G’ values to a non-parametrically estimated null distribution, with the assumption that the distribution of the G’ is approximately log-normal (See methods for details). To identify if genetic differentiations correlate with the detected QTLs, population differentiation (*F_st_*) between dead and alive pools was calculated at a sliding window and gene-wise. To identify consistent changes in allele frequencies for several replicates, we performed the Cochran-Mantel-Haenszel (CMH) test for all possible comparisons independently for each permethrin family. Our BSA approach successfully identified multiple QTLs associated with both DDT and permethrin resistance, demonstrating the method’s effectiveness in detecting resistance-associated genomic regions while identifying SNPs within these regions (Table 1).

**Table 1.** Number of QTLs identified in each BSA analysis, along with the number of genes, total significant SNPs and Nonsynonymous SNPs located within detected QTLs.

| Family | Comparison | Number of QTLs | Number of genes in QTLs | Total number of significant SNPs | Missense variant SNPs |
| --- | --- | --- | --- | --- | --- |
| DDT-A | AL-vs-DE | 4 | 1,013 | 172,103 | 2,512 |
| DDT-B | AL-vs-DE | No QTLs detected |  |  |  |
| Perm-A | AL25-vs-DE3 | 2 | 811 | 41,023 | 1,263 |
| Perm-A | AL25-vs-DE8 | No QTLs detected |  |  |  |
| Perm-A | AL30-vs-DE3 | 3 | 3,458 | 206,377 | 5,872 |
| Perm-A | AL30-vs-DE8 | 2 | 1,640 | 169,970 | 4,002 |
| Perm-B | AL60-vs-DE20 | 7 | 4,617 | 301,191 | 8,633 |
| Perm-B | AL60-vs-DE15 | 2 | 1,736 | 120,675 | 3,165 |
| Prem-B | AL75-vs-DE20 | No QTLs detected |  |  |  |
| Prem-B | AL75-vs-DE15 | No QTLs detected |  |  |  |

### BSA analysis using DDT-resistant crosses revealed a major resistance locus on chromosome 3R overlapping the GST epsilon cluster

BSA analysis performed on pools from the progeny of the crosses established between the DDT-resistant ZAN/U strain and the FAST5 susceptible strain detected a total of four significant quantitative trait loci (QTL) in one of the DDT crosses (DDT-A), but no QTL were detected in the DDT-B family (Table 1, Table S2). Performing several tests to determine the significance of BSA QTLs was essential since the commonly used delta SNP-index test didn’t detect any significance (Figure 2C). The most prominent signal was detected on chromosome 3R, where the G-prime statistic peaked at 4.05, substantially exceeding the significance threshold. This major resistance locus spanned 11.2 Mb from position 25.3-36.5 Mb on chromosome 3R, with the maximum G-prime value occurring at position 28.3 Mb (Figure 2D) (Table S2 and S3). Notably, this region contains the *GSTe* gene cluster, including *GSTe1* to *GSTe8* located between 28.58-28.6, cytochrome P450 genes (*CYP9M1* and *CYP9M2*), *GSTu3*, and a cluster of eight ABC transporter genes from the G subfamily (*ABCG8*, *ABCG10*, *ABCG11*, *ABCG16*, *ABCG17*, *ABCG18*, *ABCG19*, and *ABCG20*) located between positions 33.9-34.0 Mb (Table S4). This major QTL (*qtl1*) overlaps with the *rtd1*-resistant QTL detected previously by Ranson et al. (2000) on the same cross using microsatellite-based QTL mapping ^25^.

**Figure 2.**
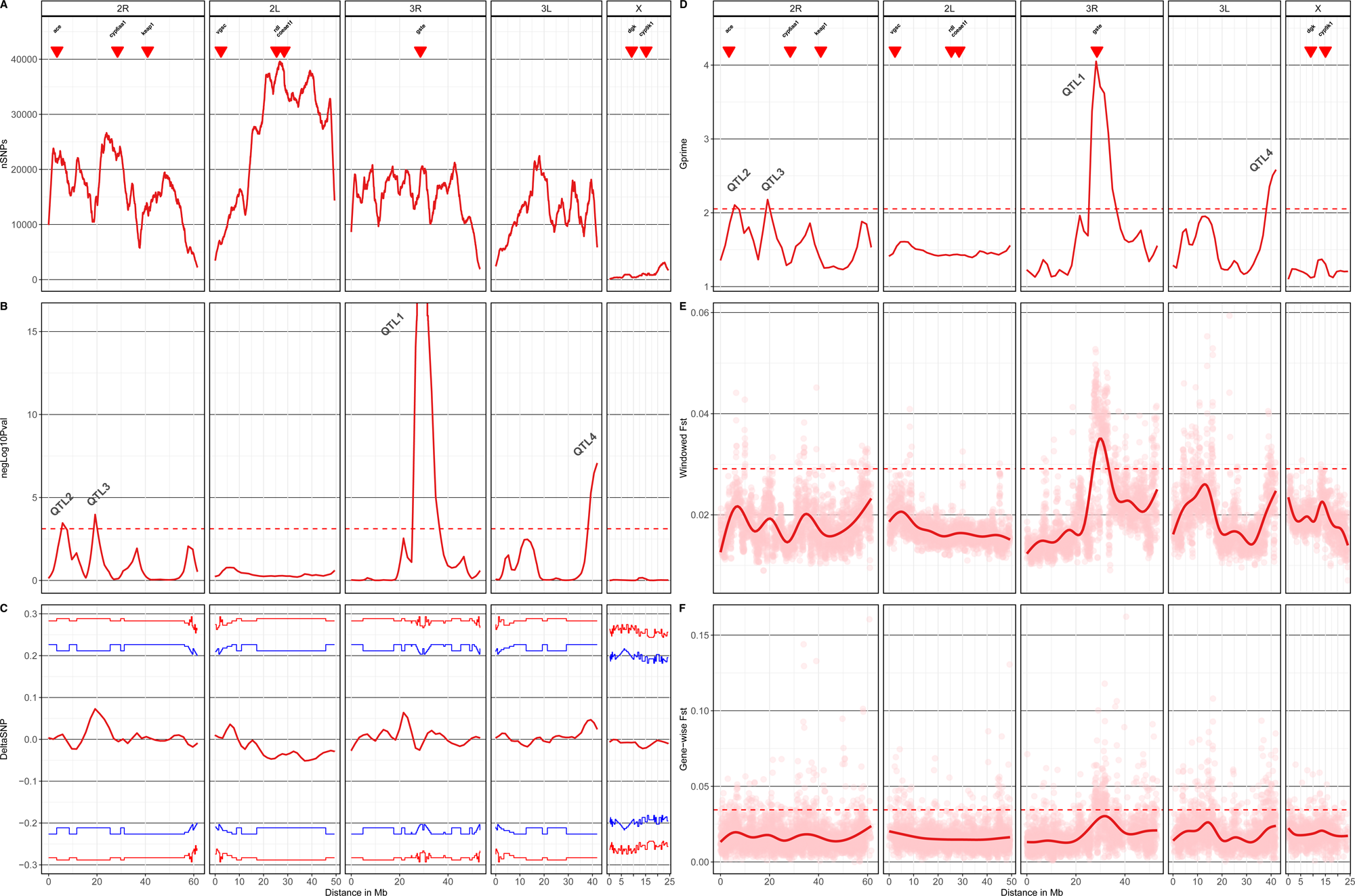
BSA in the DDT-A family identified a significant peak at the GST epsilon. Pool sequencing obtained a sufficient SNP density genome-wide. **(A)** SNP density plot in a window size of 3 Mb across all chromosomes, highlighting the number of SNPs detected in each window across the genome. High SNP density was detected across the genome, except around centromeric and telomeric regions. **(B)** Negative log10 p-value and Benjamini-Hochberg adjusted p-values were calculated on the (y-axis), showing the prominent QTL peak (*qtl3*) on the 3R chromosome; red-dotted line indicates the threshold at a genome-wide FDR of q = 0.01. The negative log10 p-value at the 3R corresponding to *qtl3* peak is equal to infinity because the p-value is effectively equal to zero, indicating a highly significant result. It was essential to perform several tests to detect BSA significance since the commonly used delta SNP-index test (**C**), highlighting the difference in SNP-index between alive and dead pool, did not detect any significance. On the other hand, G’ value calculated at a window size of 3 Mb (**D**) highlights the major QTL (*qtl3*) located at 3R spanning 25.3-36.5 Mb, with a maximum G-prime value of 4.05 at position 28.3 Mb. The peak with the strongest signal overlaps with the *GSTe* cluster region highlighted by the red arrow. Red arrows indicate genomic regions of insecticide resistance genes commonly found to be under selection in *An. gambiae* field collections. **(E)** Windowed *F_ST_* analysis at a window size of 1 Mb showing genetic differentiation between resistant and susceptible pools across the genome. Elevated *F_ST_* values beyond the 0.95 quantile, highlighted by a dotted red line cluster around QTL regions, corroborate the BSA findings. A notable differentiation is observed around the QTL corresponding to the *GSTe* cluster **(F).** Gene-wise *F_ST_* analysis displaying differentiation at the individual gene level across the genome, with notable differentiation in genes located around the major QTL region, with *F_ST_* values beyond the 0.95 quantile.

Additional significant QTLs, but with less effect compared to the strong signal detected at the 3R, were observed on chromosome 2R, with two regions spanning 5.4-7.3 Mb (*qtl2*) and 18.7-20.1 Mb (*qtl3*) showing G-prime values of 2.10 and 2.18, respectively. No major detoxification enzymes were detected in these 2R chromosome QTLs. A notable QTL (*qtl4*) was also detected on chromosome 3L, spanning 3.8 Mb from position 38.1-42.0 Mb with a maximum G-prime of 2.58. This 3L region harbours a cluster of cytochromes P450 genes including *CYP9J3*, *CYP9J4*, *CYP9J5*, *CYP9L1*, *CYP9L2*, and *CYP9L3*, as well as two additional ABC transporter genes (*ABCA5* and *ABCF2*) (Figure 2) (Table S3). No significant QTL was detected at 2L corresponding to *rtd2*, despite using individuals from the same cross for the bulk sequencing ^25^.

Windowed and gene-wise population differentiation analyses corroborated the QTLs. The windowed *F_ST_* and Gene-wise *F_S_*_T_ analysis revealed elevated differentiation around all QTL regions identified by the BSA analysis, beyond the 0.95 quantile of *F_S_*_T_ values genome-wide. However, a notable concentration of genetic differentiation is observed in the 3R region containing the *GSTe* cluster (Figure 2E and F). In contrast to the strong associations detected in family DDT-A, analysis of family DDT-B did not reveal any genomic regions exceeding the significance threshold (Figure S1). This difference between families suggests potential heterogeneity in resistance mechanisms or genetic backgrounds influencing the strength of association signals and highlights the importance of analysing multiple families in genetic mapping studies of insecticide resistance.

### BSA identifies extensive promoter and nonsynonymous SNPs within the GST epsilon cluster, potentially associated with resistance to DDT

Next-generation sequencing of DNA pools coupled with BSA allows the detection of significant SNPs in QTL regions and highlights the SNP-index difference between the pools (Table S5). BSA analysis in the DDT-A family identified, in total, 172,105 BSA-significant SNPs in all QTL regions (Table 1). Most of these SNPs were in noncoding regions; however, 2,704 SNPs were identified to cause missense mutations (Table S3). Fine-scale analysis of the major 3R QTL revealed numerous SNPs within the *GSTe* cluster, with a significant difference in SNP-index between the resistant and susceptible pools (Figure 2E). A total of 775 significant SNPs were identified across *GSTe* genes (*GSTe1*-*GSTe8*) and *GSTu3*, of which 22 SNPs were identified to be nonsynonymous SNPs (Figure 3, Table S5).

**Figure 3.**
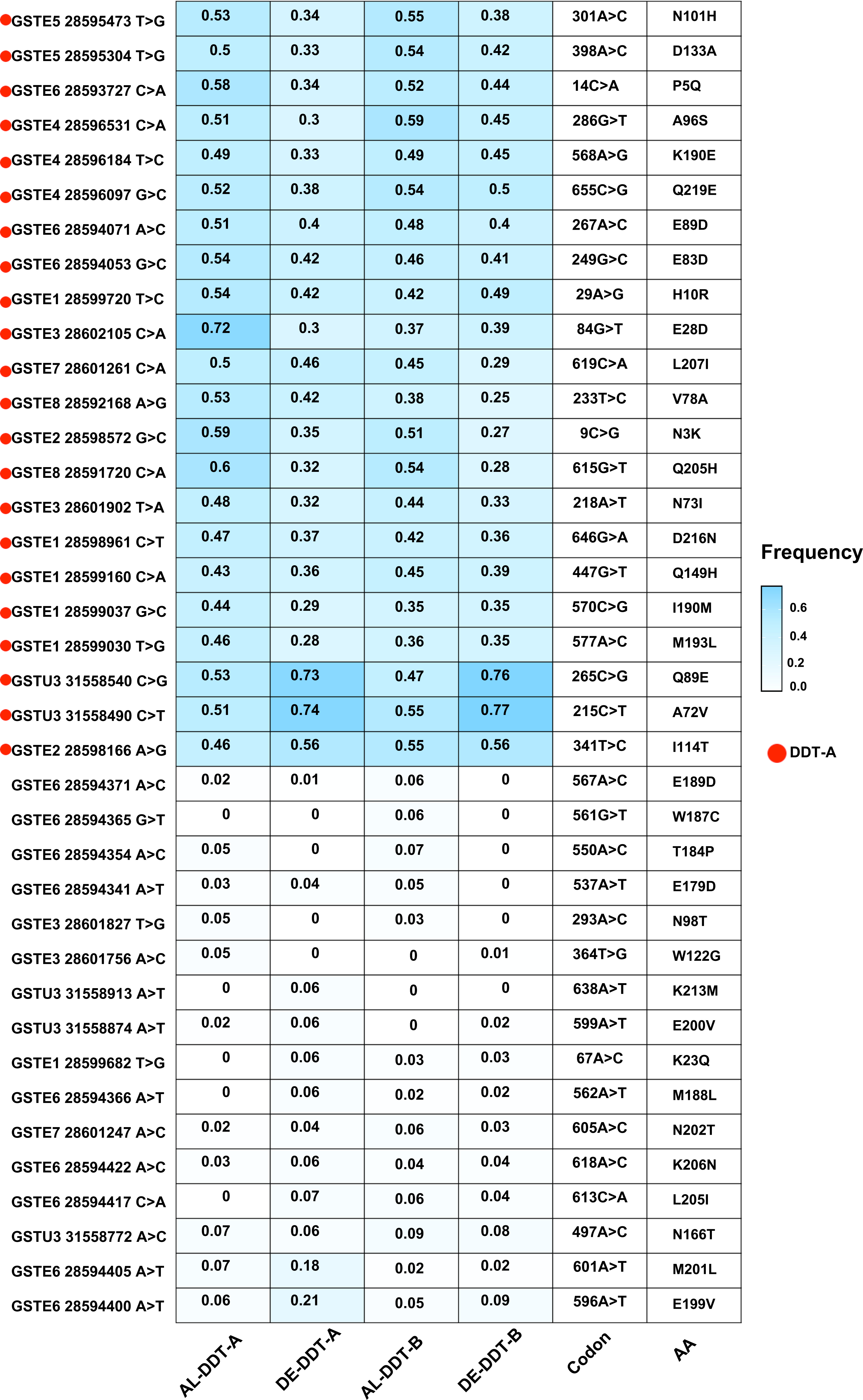
A heatmap of nonsynonymous SNPs in the *GSTe* cluster was identified using BSA. A heatmap summarising nonsynonymous SNPs in *GSTe* genes within the 3R QTL region identified using BSA, including gene names, chromosomal positions, codon and amino acid changes, and the SNP-index across all pools from the DDT-A and DDT-B crosses. Red dots highlight nonsynonymous SNPs identified as significant in the BSA analysis of the DDT-A family, whereas no significant regions were detected in the DDT-B family.

In the DDT-resistant ZAN/U strain, transcription of *GSTe2* and *GSTe3* was reported to be significantly higher compared to susceptible strains, particularly *GSTe2* and *GSTe3* ^36^. Several polymorphisms in the promoters of *GSTe2* and *GSTe3* in the DDT-resistant ZAN/U strain have been previously reported to be potentially responsible for increased promoter activity ^36,37^. Therefore, we investigated polymorphism in the 1300 bp region immediately upstream of *GSTe3* (the first gene on the reverse strand of the *GSTe* epsilon), and the 352 bp intergenic region between the stop codon of *GSTe1* and the translation start codon of *GSTe2*. BSA analysis detected a total of six SNPs with a significant SNP-index difference in the promoter region of *GSTe2,* with a significantly high SNP-index in the resistant pool (Figure 4 and Figure 5). Previously, luciferase assay using progressive deletion constructs of the *GSTe2* promoter identified that sequence polymorphisms between −301 and −255 bp were important in determining transcription rates ^37^. In this region, we detected a T>C substitution -292 upstream of the transcription start site and a T>G substitution at -283 with a significantly higher SNP-index in the resistant pool compared to the susceptible. Additional polymorphism was detected in the rest of the promoter region, including G>A -249, T>A -243, T>G -213 and A>C -185 (Figure 4). In the *GSTe3* promoter, the region between −1292 and −1039 from the transcription start site was previously determined to be essential for increased promoter activity ^37^.

**Figure 4.**
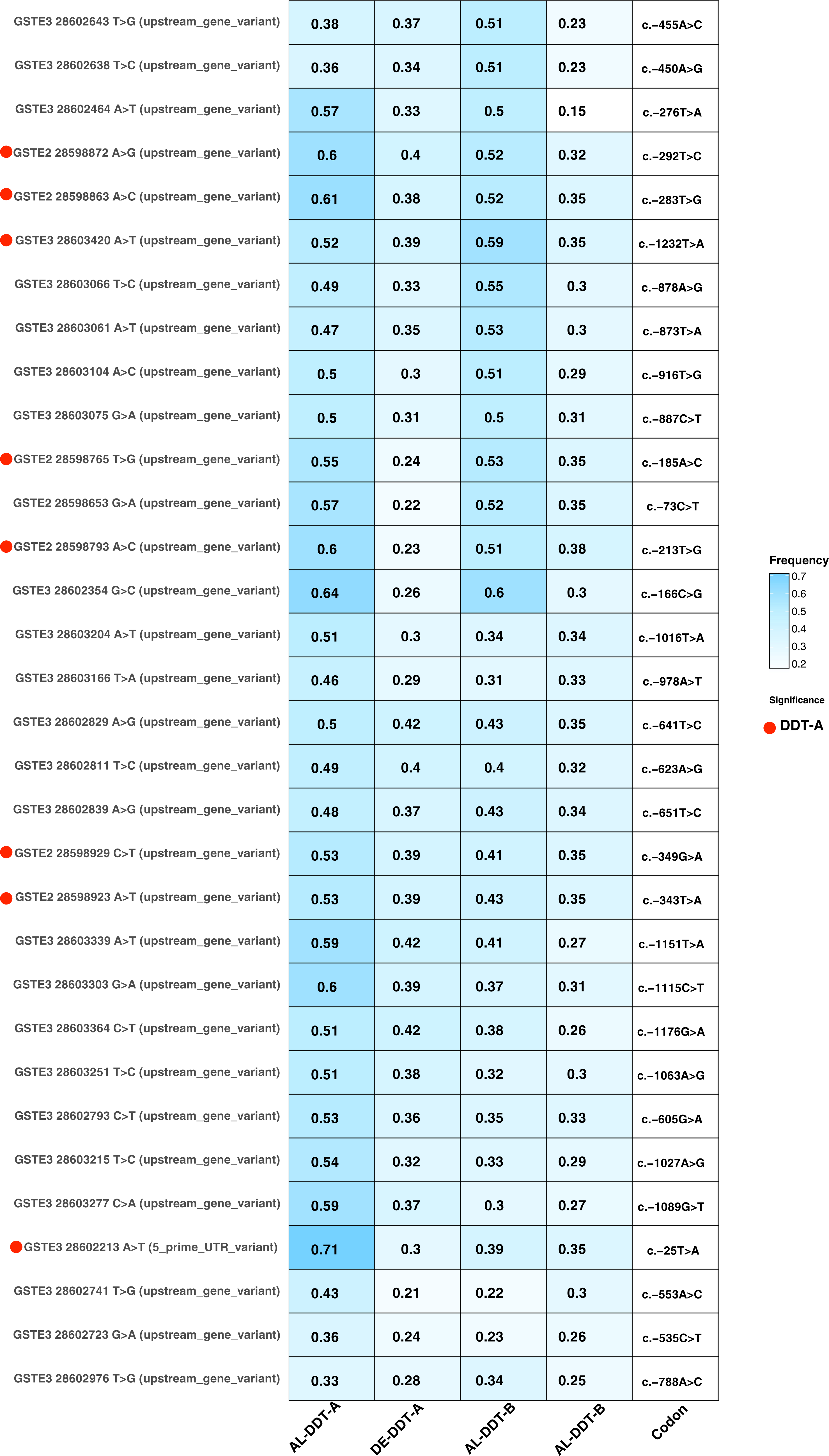
Polymorphisms in the promoter region of the *GSTe2* and *GSTe3* genes. A heatmap summarising SNPs in the promoter region of *GSTe2* and *GSTe3* genes, including gene names, chromosomal positions, the distance from the transcription start site and the SNP-index across all pools from the DDT-A and DDT-B crosses. Red dots highlight SNPs identified as significant in the BSA analysis of the DDT-A family.

**Figure 5.**
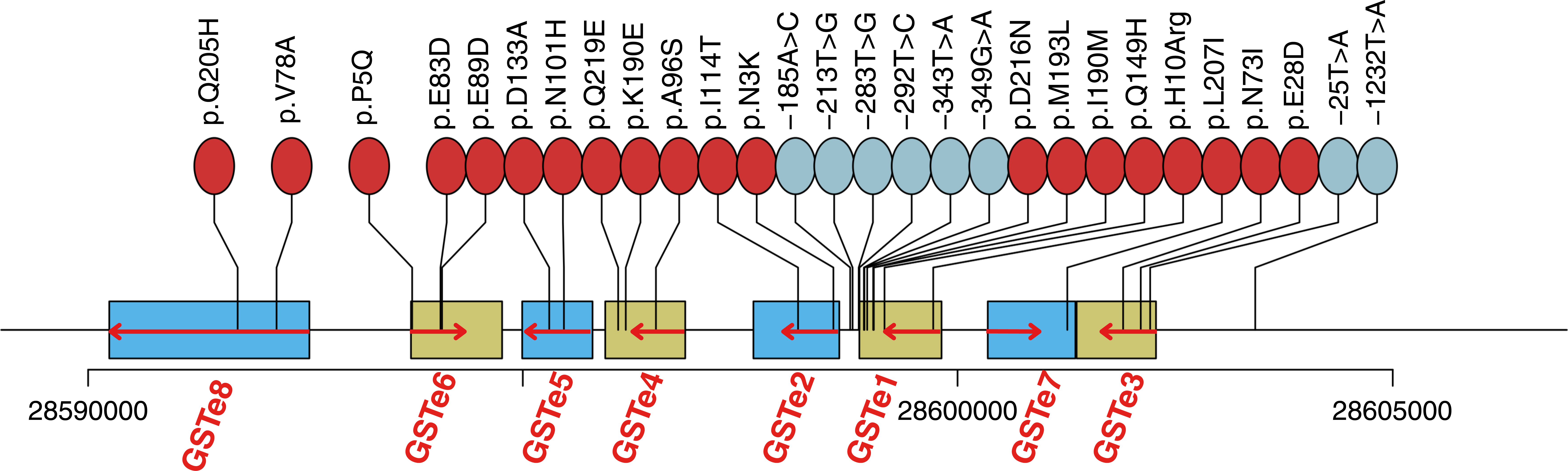
The position of *GSTe* BSA-significant SNPs. Nonsynonymous SNPs (Red) and significant SNPs in the promoter region of *GSTe2* and *GSTe3* (Blue) relative to *GSTe* genes. The direction of GSTe gene transcription is represented with a red arrow highlighting *GSTe* genes on the reverse strand.

In this region, we identified a T>A substitution at nucleotide position -1232 from the *GSTe3* transcription start site, with a significantly higher SNP-index in the resistant pool compared to the susceptible (Figure 4). In addition, a SNP in the 5’ UTR region of *GSTe3,* T>A substitution at the -25 nucleotide position upstream TSS had a significantly high SNP-index (0.71) in the resistant pool. These promoter SNPs show a significantly higher SNP-index in the resistant pool compared to the susceptible pool, highlighting their role in potentially driving the overexpression of *GSTe2* and *GSTe3* in resistant pools.

Additionally, BSA analysis in DDT-A family discovered nonsynonymous SNPs with a significant difference in SNP-index between resistant and susceptible pools distributed across all *GSTe* paralogs: *GSTe1* (4 variants: E28D, N73I, A72V, M193L), *GSTe2* (2 variants: I114T, N3K), *GSTe3* (3 variants: N73I, E28D, Q89E), *GSTe4* (3 variants: Q219E, K190E, A96S), *GSTe5* (2 variants: N101H, P5Q), *GSTe6* (3 variants: E199V, E83D, E89D), *GSTe7* (1 variant: L207I), and *GSTe8* (2 variants: V78A, D216N). Additionally, nonsynonymous SNPs were identified in GSTU3 (2 variants: A72V, Q89E), which colocalise with the *GSTe* cluster in the same 3R candidate region (Figure 3). Notably, established resistance mutations showed varied patterns in our crosses. The well-characterised *GSTe2* I114T resistant mutation showed moderate frequencies in DDT-resistant pools, with a slightly higher frequency in the dead pool from DDT-A (0.56) compared to the alive pool (0.46). The *GSTe2* I114T is a known DDT resistance marker predominantly reported in West Africa (Benin, Burkina Faso, Ghana, and Gambia), with a higher frequency in *An. coluzzii* ^38–41^. In contrast, this mutation is rare or absent in East Africa, as confirmed by MalariaGen allele frequency showing no detection in East African *An. gambiae* samples (Figure 7). Given the geographic distribution of this mutation and the slightly low frequency in alive compared to dead mosquitoes in our DDT-A pools, we consider it unlikely that this nonsynonymous SNP contributes to the resistance phenotype observed in our study populations.

### BSA analysis using historical permethrin crosses identified a major QTL harbouring *vgsc* gene

To identify QTLs associated with permethrin resistance using BSA, DNA pools were constructed using intercross offspring from crossing families Perm-A (F4 offspring) and Perm-B (F5 offspring). Mosquitoes from the original cross were phenotyped based on permethrin exposure duration in a time-course approach. In total, four pools were constructed per family, representing extreme phenotypes: long-duration survivors and short-duration mortalities (see methods for details). BSA was conducted between pools of long-duration survivors and short-duration survivors, comprising four different BSA analyses per family. Bulk segregant analysis of permethrin resistance in families Perm-A and Perm-B consistently identified two major genomic regions at chromosome 2L (approximately, 8.3 Kb – 3.7 Mb) and chromosome 2R (approximately, 47.5 – 60.2 Mb) with G-prime values reaching 20 (Table 1 and Figure 6). Each BSA comparison in both families discovered a different number of additional QTLs with varying lengths, with only the QTL in the centromeric region of chromosome 2 consistently detected in all permutations. We performed the Cochran-Mantel-Haenszel (CMH) test for all possible comparisons independently for each family to identify consistent changes in allele frequencies from the different pool comparisons. A consistent difference in allele frequency between dead and live pools in the centromeric region of chromosome 2, exceeding the threshold for Bonferroni correction, was identified (Figure 6B and C).

**Figure 6.**
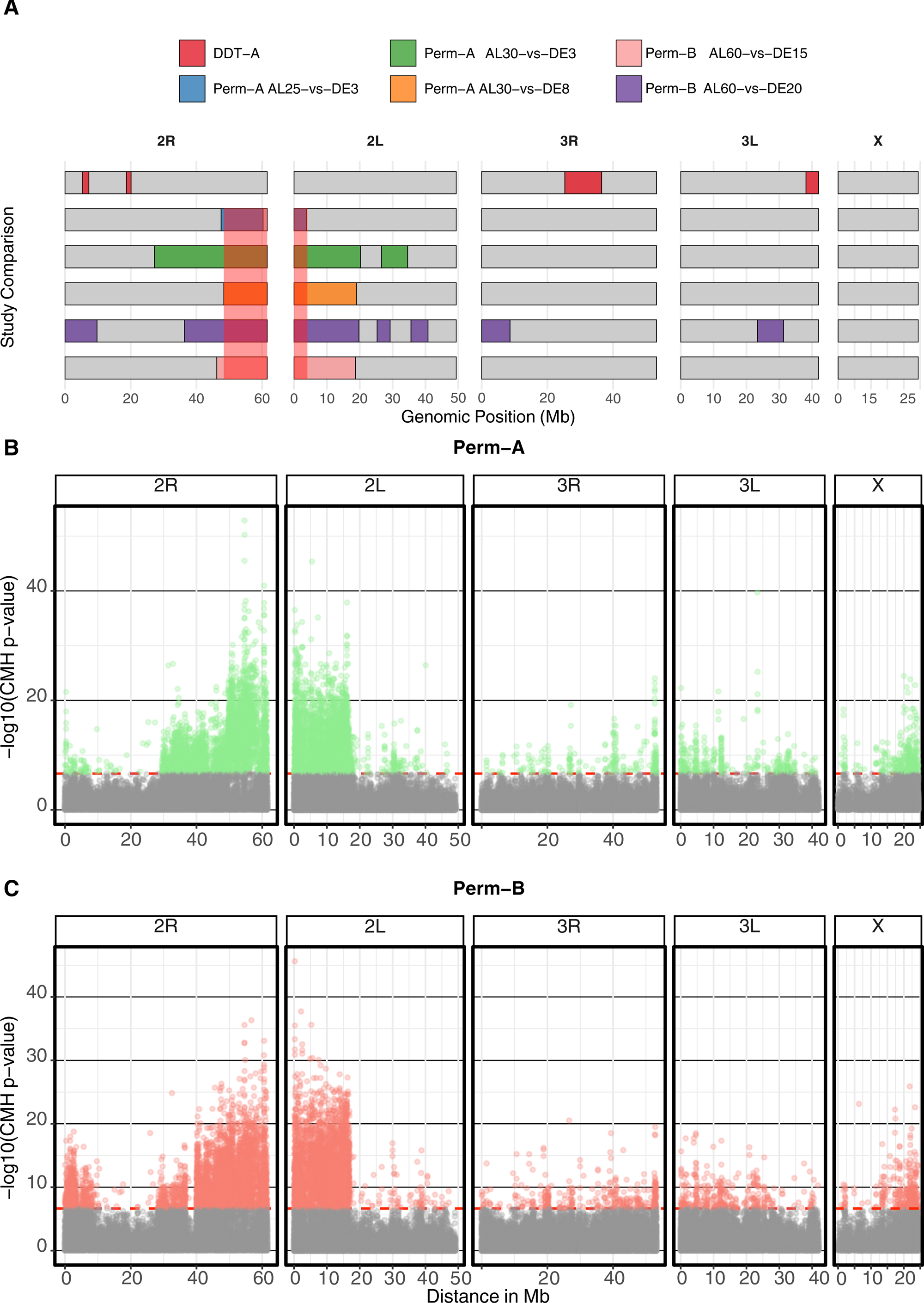
Quantitative trait loci (QTL) identified using BSA analysis and CMH test in permethrin crosses. **(A**) Detailed view of QTL regions discovered in this study, with different colour bars for each pairwise comparison where significance was recorded. The red box on 2R and 2L highlights the QTL regions discovered in consensus across different comparisons. (**B**) CHM test in Perm-A family. (**C**) CHM test in Perm-B family.

The detected QTL on chromosome 2L harbours the voltage-gated sodium channel gene (*vgsc*) at position 2.36 - 2.43 Mb, the primary target of pyrethroid insecticides and a well-established resistance gene in *An. gambiae*. This region corresponds to the *rtp1* locus conferring resistance to permethrin, identified in the same cross by microsatellite QTL mapping in ^28^. The RSP-ST strain had already been confirmed to contain a leucine to serine substitution at position L1014S according to *Musca domestica* codon numbering corresponding to L995S codon in *An. gambiae* transcript AGAP004707-RD ^9^. The L995S *kdr* mutation (2L:2,422,651 T>C) is present at a higher frequency in permethrin survival pools in both families (Figure S7), although only significant in the Perm-A family. The L995S *kdr* mutation was also observed at a very low frequency (0.02 – 0.13) in DNA pools from the DDT families, illustrating that the *kdr* mutation is nearly absent in DDT families and metabolic detoxification is the primary mechanism of resistance (Figure S7). Another *vgsc* mutation, M490I (2L:2400071 G>A), shows significant associations in multiple permethrin BSA comparisons. Although the M490I mutation has not been established as a knockdown-resistant mutation (*kdr*), recently it was demonstrated to be potentially under selection in the Kenyan population ^8^. The QTL region discovered from BSA in the 2L chromosomes also contains carboxylesterase genes (*COEBE1C-4C*) and prophenoloxidase genes (*PPO3*, *PPO4*, *PPO6*, *PPO7*, *PPO8*, *PPO9*), though these may represent linkage with the *vgsc* locus rather than independent selection ^38^.

The QTL on chromosome 2R 47.5 – 60.2 Mb contains multiple detoxification genes, including a major glutathione S-transferase delta cluster (*GSTD1-12*), glutathione peroxidases (*GPXH1*, *GPXH3*), and ABC transporter genes (*ABCG4*, *ABCC4*, *ABCC6*). Genes from the GST delta cluster were previously reported to be overexpressed across the *An. gambiae* complex in multiple locations across Africa; however, no previously recorded signature of selection was detected at these genes ^42^. Given the QTL strong signal at the well-characterised *vgsc* locus, the 2R signal may reflect linkage disequilibrium rather than independent metabolic resistance mechanisms. A QTL was detected on 3R in the Perm-B family, corresponding to the previously reported *rtp2* (Ranson et al., 2004), when BSA was conducted between the alive pool after 60 minutes of exposure and the dead pool after 20 minutes of exposure (Figure 6A and Table S14) ^28^. This QTL contains several *CYP6* genes, including *CYP6Z1*, which was reported to be overexpressed in adults from the resistant RSP-ST strain compared to insecticide-susceptible adults ^31^. We did not t detect other QTLs in 3R that corresponded to the previously detected *rtp3* QTL ^28^.

A complete list of genes identified in candidate regions for each BSA analysis and SNPs with significant SNP-index difference between the two pools for each family where QTLs were detected are provided in tables S6-S15. These findings suggest that permethrin resistance in the two intercrossed families used in the BSA analysis is primarily driven by target-site modifications at the *vgsc* locus.

### GST epsilon mutations show variable geographic distribution patterns in East African *An. gambiae* populations

To contextualise our BSA findings within broader signatures of genetic selection, we examined the distribution of BSA significant nonsynonymous and promoter mutations in the *GSTe* epsilon across East African *An. gambiae* populations using data from the Malaria Genomic Epidemiology Network (MalariaGEN) ^43^. We focused on a subset of MalariaGEN *An. gambiae* specimens representing diverse ecological zones and insecticide exposure histories collected from East African countries (Kenya, Tanzania, and Uganda), since the resistant laboratory strain (ZAN/U) was originally isolated from the Zanzibar population. In total, we analysed a subset of 2,789 *An. gambiae* mosquitoes collected from 405 sites between 2000 and 2019. Most of the samples came from Uganda (2,432 mosquitoes from 362 sites), followed by Tanzania (301 mosquitoes from 35 sites) and Kenya (56 mosquitoes from 7 sites). Collection timing varied by country, with the Kenya samples spanning the longest period (2000-2019) (Figure 7, Table S16).

**Figure 7.**
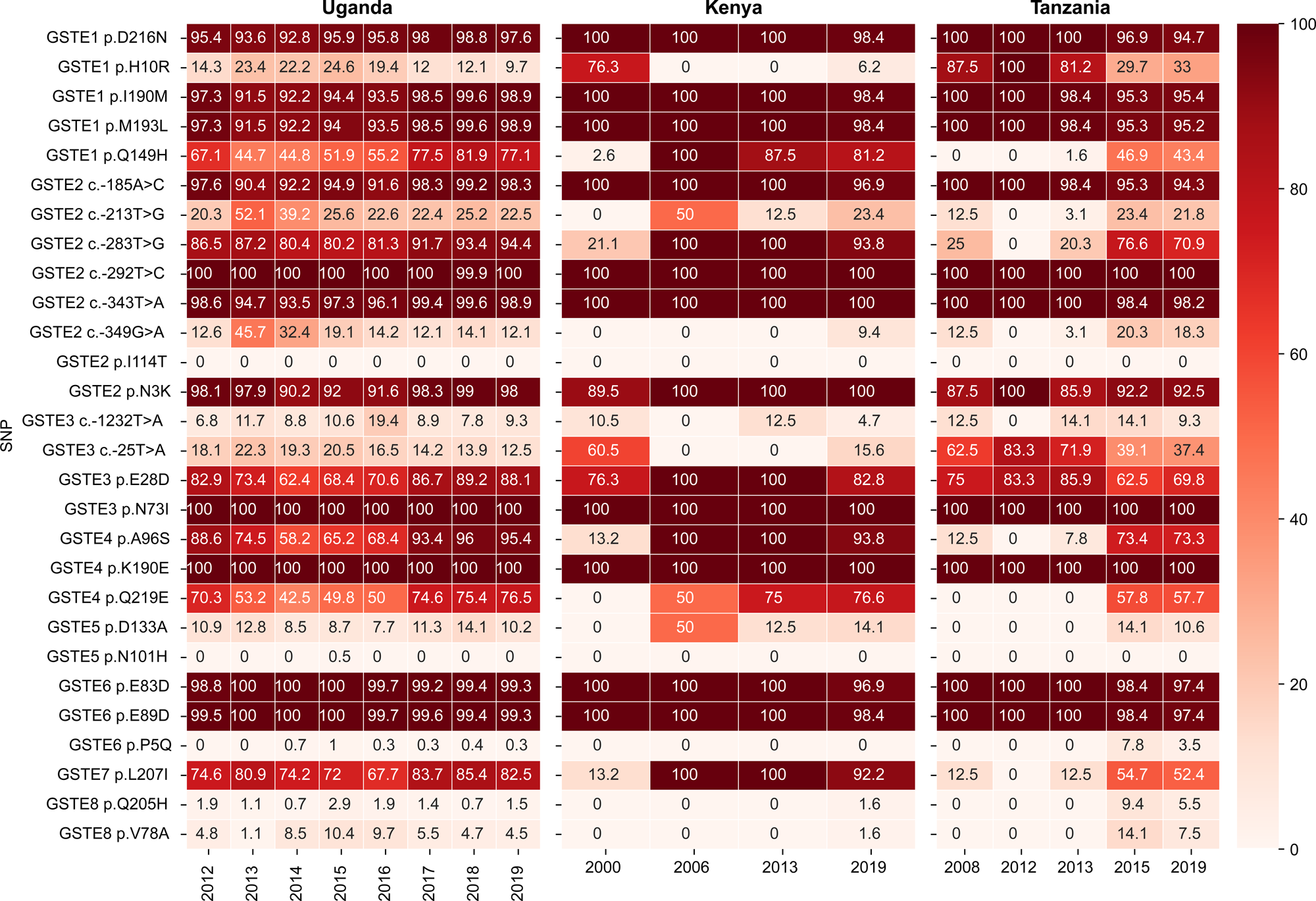
Allele frequency of GST epsilon SNPs in all *An. gambiae* East African samples per year from the MalariaGEN network. Nonsynonymous SNPs and promoter SNPs identified as significant from the BSA analysis were investigated in the *An. gambiae* samples from East African countries, including Uganda, Kenya and Tanzania. Nonsynonymous SNPs causing an amino acid substitution are denoted with a ‘p.’ prefix, while SNPs causing nucleotide-level substitution at the promoter region are denoted with a ‘c.’ prefix. The allele frequency of SNPs (y-axis) is plotted in a heatmap for each collection year (x-axis) in each country.

Analysis of the 20 nonsynonymous SNPs within the *GSTe* cluster and SNPs upstream of *GSTe2* and *GSTe3* discovered from BSA revealed distinct evolutionary trajectories across East African *An. gambiae* populations, possibly reflecting differential selection pressures and the temporal dynamics of metabolic resistance evolution. A total of eight nonsynonymous variants in *GSTe1*, *GSTe3*, *GSTe4*, and *GSTe6* are nearly fixed (>90%) across Uganda, Kenya, and Tanzania throughout 2000 to 2019, indicating these mutations may confer advantages under insecticide pressure that were already established during the intensifying DDT vector control era (Figure 7). We also identified three nearly fixed BSA-significant variants in the promoter region upstream of *GSTe2* in East African samples, located in the intergenic space between *GSTe1* and *GSTe2*. These variants are nearly fixed with frequencies > 90% consistently from 2000 to 2019 across all three East African countries. Given their association with the resistant promoter copy, these variants may represent beneficial mutations that have been strongly selected for their ability to enhance *GSTe2* expression under intensive insecticide pressure, warranting future laboratory validation.

Interestingly, several nonsynonymous variants demonstrated clear progression towards high frequency from near absence, especially in Kenya and Tanzania, suggesting ongoing adaptive evolution. For example, *GSTe7* L207I increased from 67.7% to 85.4% in Uganda, while rising dramatically from 13.2% (2000) to 100% (2019) in Kenya and from 12% (2008) to 52.4% (2019) in Tanzania, likely reflecting intensifying selection pressure. Similarly, *GSTe1* Q149H emerged from a complete absence to 81.2-100% in recent Kenyan collections and 43% in the most recent collection from Tanzania. *GSTe4* Q219E emerged from a complete absence in Tanzania and Kenya to a frequency of 76.6% and 57.7%, respectively, in the most recent collection in 2019. A similar pattern was observed for the *GSTe2* promoter variant at -283 T>G, where the frequency increased to near fixation in the most recent collections. The increase in frequency of the nonsynonymous and promoter SNPs from near absence in the earliest collection from the MalariaGEN dataset highlights continued evolution and ongoing refinement of the metabolic machinery to optimise detoxification efficiency and maintain fitness under sustained insecticide pressure.

## Discussion

This study demonstrates that Genetic crosses followed by Bulk Segregant Analysis (BSA) offers a robust method to identify insecticide resistance loci in *An. gambiae* ^44,45^. Next-generation sequencing, coupled with BSA, identified several of the same QTLs previously detected using microsatellite markers. The major DDT QTL (*qtl3*) on the 3R chromosome co-localised with the *GSTe* cluster. Amino acid substitutions and promoter polymorphism with a significant SNP-index difference between resistant and susceptible pools were identified, which were present when the *ZAN/U* population was colonised in the 1980s. Many of these historically derived mutations show signatures of long-term selection pressure across Eastern African countries. DDT was phased out for malaria control early in the 20^th^ century, following the Stockholm Convention banning the use of this insecticide ^46^. Despite this, stockpiles of DDT are still known to exist in Tanzania, and unlicensed use in agriculture persists, potentially contributing to the selection for polymorphisms in the GSTe cluster that increase *Anopheles*’ ability to detoxify this insecticide ^47^.

The BSA-significant polymorphisms identified within the *GSTe* cluster may represent some of the earliest resistance alleles to evolve within this critical detoxification cluster. When these SNPs were investigated using East African *An. gambiae* field samples from the MalariaGEN network collected > 20 years after the ZAN/U strain was colonised, it was revealed that many of the historically derived mutations identified from BSA analysis have persisted, and several have increased in frequency in Eastern Africa during the 21^st^ century. Whether this is due to continued DDT exposure or indicative of alternative selection pressures on this gene cluster is unknown. Increased GST activity has been previously associated with pyrethroid resistance, and glutathione conjugation may be an important secondary detoxification mechanism to counter the effects of pyrethroid exposure in ITNs ^48^. Interestingly, GSTs have been linked to increased tolerance to PBO-pyrethroid ITNs ^49^.

Permethrin resistance in the RSP-ST strain was largely associated with target-site modifications at the *vgsc* locus on chromosome 2L. It is essential to analyse multiple crossing families when conducting BSA (and economically feasible since only two samples require sequencing to map candidate regions), since differences in genetic background and heterogeneity in resistance mechanisms may influence the number of candidate regions detected and the strength of the signal at these regions. This correspondence in the significance of the chromosome 2 QTL in permethrin resistance between the different comparisons was confirmed by the CMH test in Perm-A and in Perm-B, independently, identifying a consistent difference in allele frequency between dead and live pools beyond the genome-wide Bonferroni correction.

While BSA proved effective for identifying resistance-associated loci, several limitations should be acknowledged. Our analysis focused exclusively on SNPs, as is standard for BSA studies, because allele frequencies can be reliably estimated from read counts at biallelic positions. Indels are less tractable in pooled sequencing data due to alignment ambiguities, and these challenges are amplified when parental strains are genetically divergent, as in our crosses ^50^. Significant indel polymorphism was previously determined in the promoter of *GSTe2* and *GSTe3* between ZAN/U and susceptible strains. Therefore, sequencing the founder strains alongside pools of crossed progeny would enhance the investigation of Indels and structural variation polymorphism in the BSA detected QTL. Additionally, integrating RNA-seq with BSA would provide a more comprehensive view of resistance mechanisms by simultaneously capturing expression QTLs and differentially expressed genes, helping to resolve whether resistance arises from coding variants, regulatory changes, or both.

The persistence of historically derived resistance mutations in contemporary field populations has direct implications for current vector control strategies. The high frequencies of *GSTe* mutations across East African populations suggest that insecticide-based interventions are likely encountering populations with pre-existing metabolic resistance capacity derived from the DDT era. This “resistance debt” from historical insecticide excessive use creates a challenging baseline against which newly deployed control tools must be evaluated. Our study demonstrates the value of preserving historical biological samples for retrospective genomic analysis. The BSA methodology we developed could be readily adapted to crosses from other vector species and resistance phenotypes. The integration of BSA results with population genomic data from contemporary field samples proved particularly valuable for validating the biological relevance of BSA-derived findings. As malaria vector control programs face increasingly complex resistance challenges, understanding the evolutionary origins of resistance mechanisms is important for developing sustainable control strategies. Our work provides both methodological tools and evolutionary insights that should inform future efforts to manage insecticide resistance and protect the effectiveness of vector control interventions.

## Methodology

### Mosquito genetic crosses, DNA extraction and sequencing

This study utilised preserved mosquito samples from genetic crosses originally established in the late 1990s and maintained for over 20 years at -80 °C before genome extraction. Two separate resistance mapping experiments were conducted using different insecticide-resistant strains. In these crosses, susceptible males were mass-crossed with resistant females, then females were separated and allowed to oviposit individually. Using the F1 progeny from these crosses, single pair crosses were established by combining a single F1 male with one to five F1 females in individual mating vials. After blood feeding, the F1 females were separated and allowed to oviposit separately to establish iso-female family lines. The F2 progeny of these iso-female F1 were allowed to inter-cross for multiple generations.

For the DDT resistance QTL mapping experiment, the original crosses were established between DDT-susceptible FAST5 males and DDT-resistant ZAN/U females. The ZAN/U strain was colonised from a DDT-resistant field population collected in Zanzibar, Tanzania, in 1982, while the FAST5 strain was derived from the 4Ar/r strain through selective breeding for standard chromosome arrangements. The F2 progeny resulting from the F1 females were allowed to intercross by sibling mating to the F8 generation. F8 individuals from two families (DDT-A and DDT-B) were exposed to 4% DDT for 45 minutes following the WHO standard protocol, resulting in approximately 50% mortality. Mosquitoes were then sorted by survival status, and DNA was extracted from individual mosquitoes as described in ^45^. Briefly, to lyse mosquito tissues for DNA extraction, an individual mosquito was placed in each well of a 96-well plate with 180 μL of lysis buffer, 20 μL of proteinase K, and a metal ball bearing. Mosquitoes’ tissues were ground using a Tissue Lyser for 5 minutes at maximum frequency and then incubated in a preheated incubator at 56°C for 3 hours. Following, incubation plates were centrifuged at ∼2000 rpm in a plate centrifuge to collect condensation and liquid at the bottom of the tube. DNA extraction was conducted in 96-well plates using the DNeasy kit (Qiagen) according to the manufacturer’s recommendation, eluted in 100 μL AE buffer and stored at -20° C. DNA quality was assessed using gel electrophoresis and concentration was measured using the Quant-iT PicoGreen dsDNA Assay kit (Thermo Fisher, MA, USA). DNA from individual mosquitoes with a similar phenotype was pooled equally to construct four pools: DDT-A alive (AL-DDT-A), DDT-A dead (DE-DDT-A), DDT-B alive (AL-DDT-B), and DDT-B dead (DE-DDT-B), with each pool consisting of 80 individuals.

For the permethrin resistance QTL mapping experiment, the original crosses were conducted between permethrin-resistant RSP-ST males and susceptible FAST females. The RSP-ST strain originated from western Kenya, derived from females collected in 1992 from areas where permethrin-impregnated bednets were in use ^9^. The crosses were intercrossed to the F4 (Perm-family) and F5 (Perm-B family) generation before insecticide exposure. To maximise phenotypic segregation, mosquitoes were exposed to long and short durations of 0.75% permethrin according to WHO bioassay standards, with only long-duration survivors and short-duration mortalities retained for analysis. DNA was extracted from individual mosquitoes as described above and pooled equally to construct a total of eight pools across two families (perm-A and perm-B), with exposure times varying to capture different resistance phenotypes. For family perm-A, pools included 94 individuals surviving 25-minute exposure (AL-perm-A-25), 40 surviving 30-minute exposure (AL-perm-A-30), 64 dying after 3-minute exposure (DE-perm-A-3), and 96 dying after 8-minute exposure (DE-perm-A-8). For family perm-B, pools consisted of 96 individuals surviving 60-minute exposure (AL-perm-B-60), 48 surviving 75-minute exposure (AL-perm-B-75), 64 dying after 15-minute exposure (DE-perm-B-15), and 72 dying after 20-minute exposure (DE-perm-B-20).

Sequencing libraries were prepared by Novogene (Cambridge, United Kingdom) using standard protocols with an average insert size of 350 bp. Whole-genome sequencing was performed on the Illumina NovaSeq 6000 platform with 150 bp paired-end reads. Raw sequences used in this study were deposited in the ENA under accession number PRJEB97122.

### Pooled-template Sequencing Variant Calling

Raw sequence reads were quality-checked using FastQC (v0.11.9) ^51^ to assess base quality, adapter contamination, and overall sequence quality. Reads were aligned to the *Anopheles gambiae* PEST reference genome (GeneBank reference assembly GCA_000005575.1) using BWA-MEM2 (v2.2.1) ^32,52,53^. The resulting alignment SAM files were converted to BAM format and sorted using SAMtools (v1.13) ^54,55^. PCR duplicates were marked and removed using GATK’s (v4.6.2.0) MarkDuplicates with the REMOVE-DUPLICATES=true option. Read groups were updated to reflect sample identities using GATK’s AddOrReplaceReadGroups ^56^. After filtration of low quality-reads and PCR duplication, on average, 9.74 × 10^7^ reads were sequenced across all samples with a mapping rate of 97.8%. Read and map metrics are shown in Table S1.

Variant calling was performed using FreeBayes (v1.3.2) in parallel with the --use-best-n-alleles 4 option to reduce computational time and --pooled-continuous to accommodate for pooled samples ^57^. Variants were annotated using SnpEff (version 5.0e)^58^ and filtered using bcftools (version 1.9) to retain only high-quality, biallelic variants (QUAL > 20, GQ ≥ 99, FMT/DP ≥ 100, and no missing genotypes) ^54^. SNPs and indels were separated into individual VCF files. Variants were subsequently converted into tabular format using GATK’s VariantsToTable for downstream BSA analysis. A total of 11,013,869 variant records were called across all samples, which included 9363189 SNPs and 837393 indels. Following variants filtering, 8,137,996 SNPs (87%) and 414609 indels (50%) were retained. Per-sample variant distributions are shown in Table S1.

### Bulk Segregant Analysis (BSA) and population genetics

The BSA custom script, based on the QTLseq R package, and the complete poolseq pipeline, are available here https://github.com/alyazeeditalal/Poolseq_analysis. SNPs were filtered before running the BSA analysis based on a reference SNP-index of 0.20, minimum total depth equal to 100, maximum total depth equal to 400 and minimum sample depth of 40. The difference in SNP-index between alive and dead bulk (delta SNP-index) was calculated on a sliding window size of 3 Mb in length, while calculating the number of SNPs per window. The 3 Mb sliding window was chosen to balance signal clarity with statistical power. Smaller windows resulted in excessive noise that obscured QTL signals, while larger windows reduced resolution and potentially merged distinct peaks. Confidence intervals at 95% and 99% were estimated using the quantile from simulating the delta-SNP-index per bulk over 1000 replications. Additionally, G-statistics were calculated genome-wide and a tricube-smoothed G-statistic (G’), a smoothing method on the G value across the genome, was calculated in a sliding window of 1 Mb. G-statistics measure the goodness of fit between observed and expected values, assessing whether deviations are due to chance or associated with a phenotype of interest, and determining if the observed differences are random or biologically relevant. A p-value was determined by comparing the G’ values to a non-parametrically estimated null distribution, with the assumption that the distribution of the G’ is approximately log-normal ^35^. Negative log10 p-value and Benjamini-Hochberg adjusted p-values were calculated ^59^. Candidate regions were identified using a genome-wide false discovery rate (FDR) of 0.01. Population differentiation (*Fst*) between dead and alive pools was calculated at the SNP level, a sliding window length of 1 Mb and gene-wise using grenedalf version 0.3.0 (https://github.com/lczech/grenedalf) ^60^. To identify a consistent difference in allele frequency between dead and live pools CMH test was performed using popoolation2 (v1.201) in pools from Perm-A and in Perm-B with all possible combinations ^61^.

### Identifying SNPs from the MalariaGen API

To validate and contextualise BSA findings, we analysed contemporary field population data from the Malaria Genomic Epidemiology Network ^43^. SNP and amino acid frequency data for nonsynonymous variants within the *GSTe* cluster were accessed from East African countries (Kenya, Tanzania, Uganda, and Malawi) using the MalariaGEN API.

SNP allele frequencies were obtained using the snp-allele-frequencies function, while amino acid frequencies for nonsynonymous variants were accessed through the aa-allele-frequencies function. Geographic and temporal patterns were analysed across the available sampling timeframe (2000-2019) to understand the evolutionary dynamics of resistance alleles.

## Data availability statement

All poolseq sequences are available at the European Nucleotide Archive (ENA) under accession number PRJEB97122.

## Acknowledgements

The authors thank Simon Wagstaff and Andrew Bennett at the LSTM scientific computing team for their support in maintaining the computational cluster used to host and analyse genomic data presented in this study.

## Author Contribution Statement

T.A.Y. conceptualised and designed the study. T.A.Y. performed the formal analysis, data curation, pipeline development, and visualisation. M.M., preforemed genomic DNA extraction from historical samples. T.A.Y., N.D., and H.R. wrote the original draft. All authors contributed to the investigation and critically reviewed and edited the manuscript. H.R. supervised the project and acquired funding. N.D. validated the findings. O.A. and A.M. participated in manuscript review.

