## Supplemental figures for "The metabolic resistance blueprint: Genomic dissection of DDT resistance in historical *kdr*-free East African *Anopheles gambiae*"

**Figure S1. BSA analysis in the Perm-B family.** In contrast to the strong QTL signal detected in Perm-A, BSA analysis in Perm-B didn't detect any signal, suggesting possible heterogeneity in the resistance mechanism and the importance of analysing multiple families to detect an association signal. (**A**) SNP density plotted in a window size of 3 Mb. (**B**) negative Log10 of p-value calculated from the G-prime value by comparing to a non-parametrically estimated null distribution, with the assumption that the distribution of the G' is approximately log-normal. (**C**) Delta SNP-index highlighting the difference between the SNP-index of resistant and susceptible pools. (**D**) G-prime value calculated at a window size of 3 Mb. (**E**) $F_{st}$ was calculated on a window size of 50,000 bp and a sliding window of 25,000 bp. (F) Gene-wise $F_{st}$ is calculated $F_{st}$ using gene regions.

**Figure S2. BSA performed on Perm-A pools; alive after exposure for 25 minutes and dead after exposure for 3 minutes.**

**Figure S3. BSA on Perm-A; alive after exposure for 30 minutes and dead after exposure for 8 minutes.**

**Figure S4. BSA result from Perm-A by comparing the alive pool after exposure for 30 minutes and the dead pool after exposure for 3 minutes.**

**Figure S5. BSA results from the Perm-B family by comparing the alive pool after exposure for 60 minutes and the dead pool after exposure for 15 minutes.**

**Figure S6. BSA results from the Perm-B family by comparing the alive pool after exposure for 60 minutes and the dead pool after exposure for 20 minutes.**

**Figure S7. Nonsynonymous SNPs discovered at the voltage-gated sodium channel gene (*vgsc*).** A heatmap summarising nonsynonymous SNPs in *vgsc* gene, including gene names, chromosomal positions, and the SNP-index across all pools from the DDT-A, DDT-B, Perm-A and Perm-B crosses.

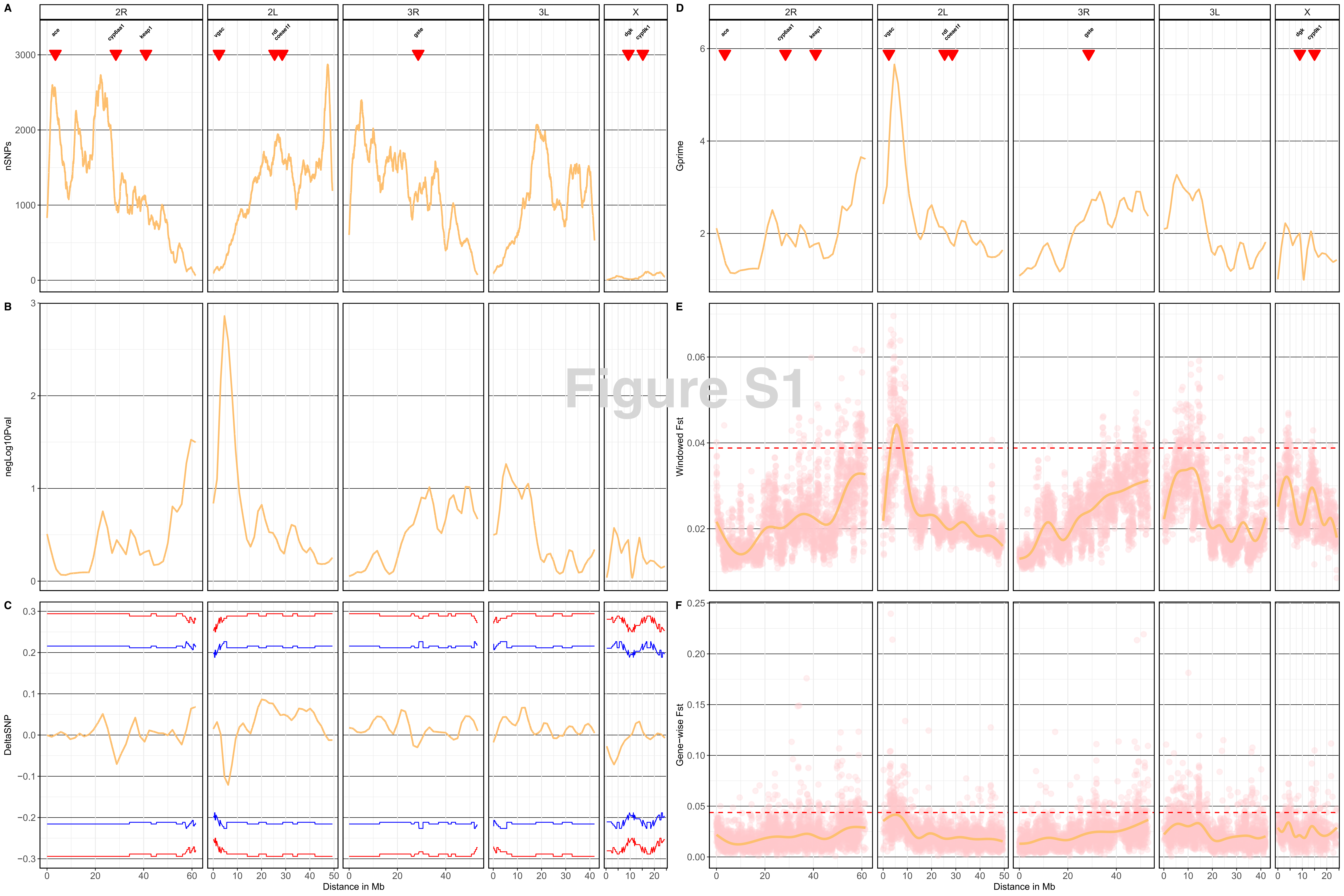

Figure S1
A
B
C
D
E
F
2R
2L
3R
3L
X
ace
cyp6aa1
keap1
vgsc
rdl
coeae1f
gste
dgk
cyp9k1
nSNPs
negLog10Pval
DeltaSNP
Gprime
Windowed Fst
Gene−wise Fst
Distance in Mb

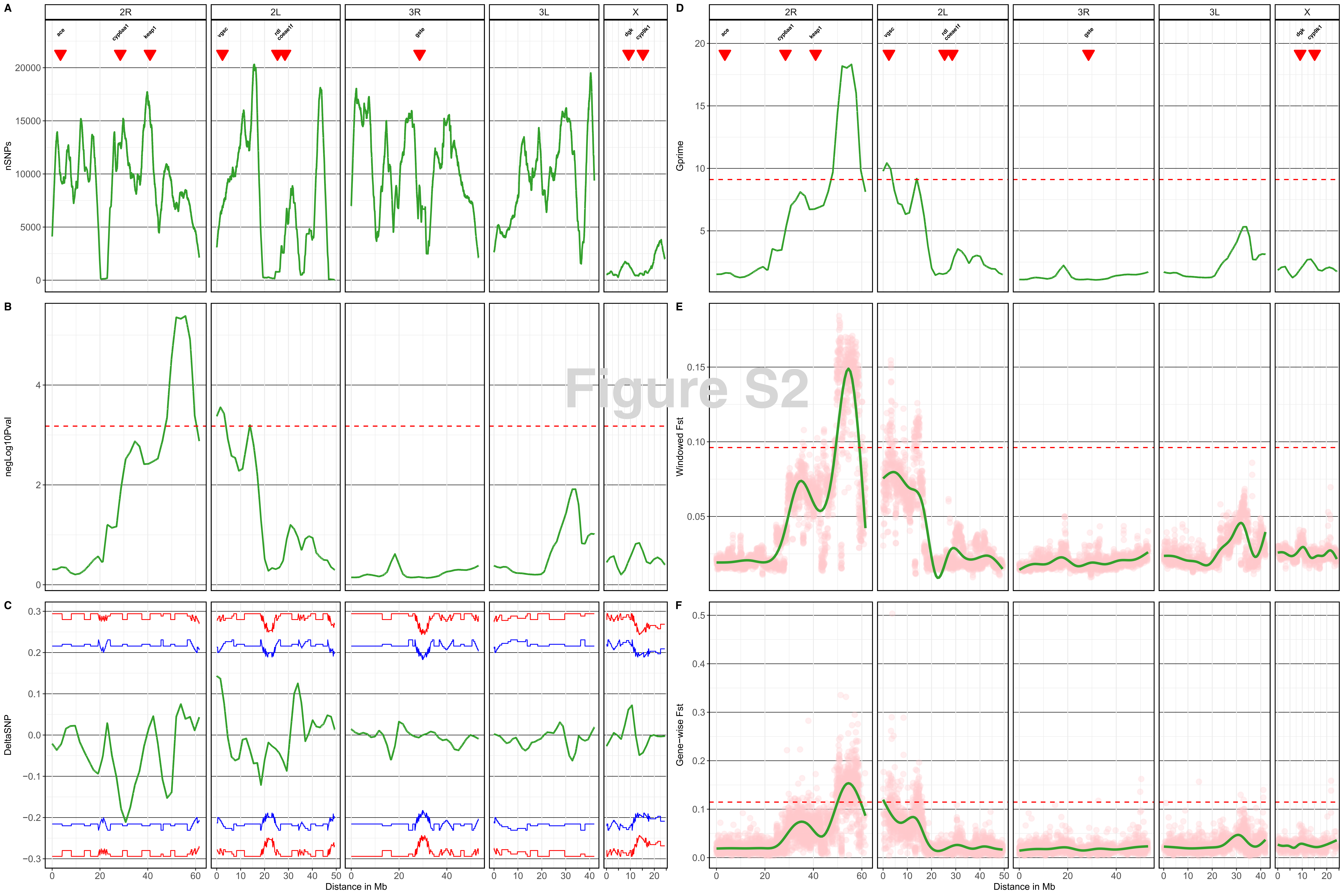

A
B
C
D
E
F
2R
2L
3R
3L
X
ace
cyp6aa1
keap1
vgsc
rdl
coeae1f
gste
dgk
cyp9k1
nSNPs
negLog10Pval
DeltaSNP
Gprime
Windowed Fst
Gene−wise Fst
Distance in Mb


Figure S2

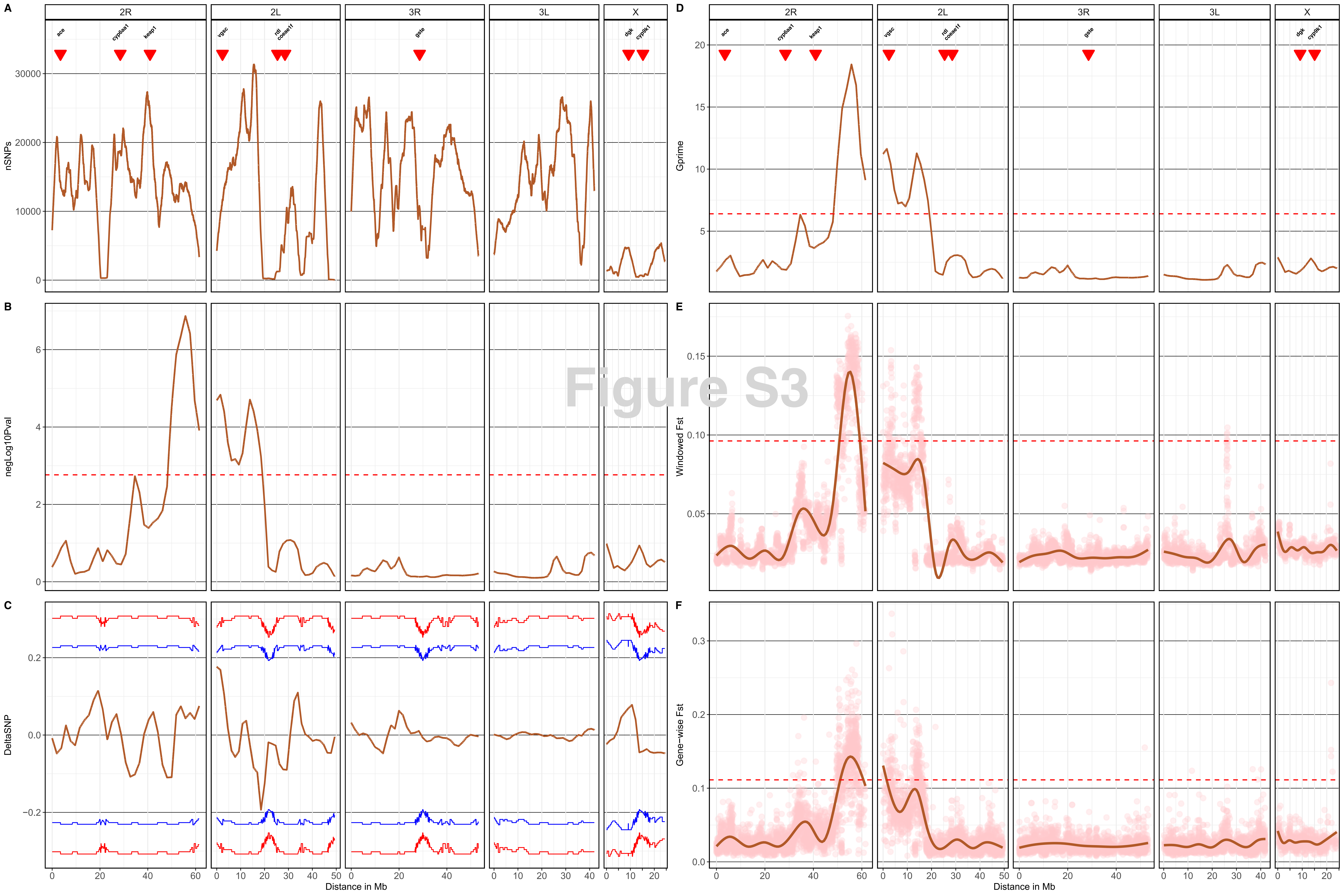

Figure S3
A
B
C
D
E
F
2R
2L
3R
3L
X
ace
cyp6aa1
keap1
vgsc
rdl
coeae1f
gste
dgk
cyp9k1
nSNPs
negLog10Pval
DeltaSNP
Gprime
Windowed Fst
Gene-wise Fst
Distance in Mb

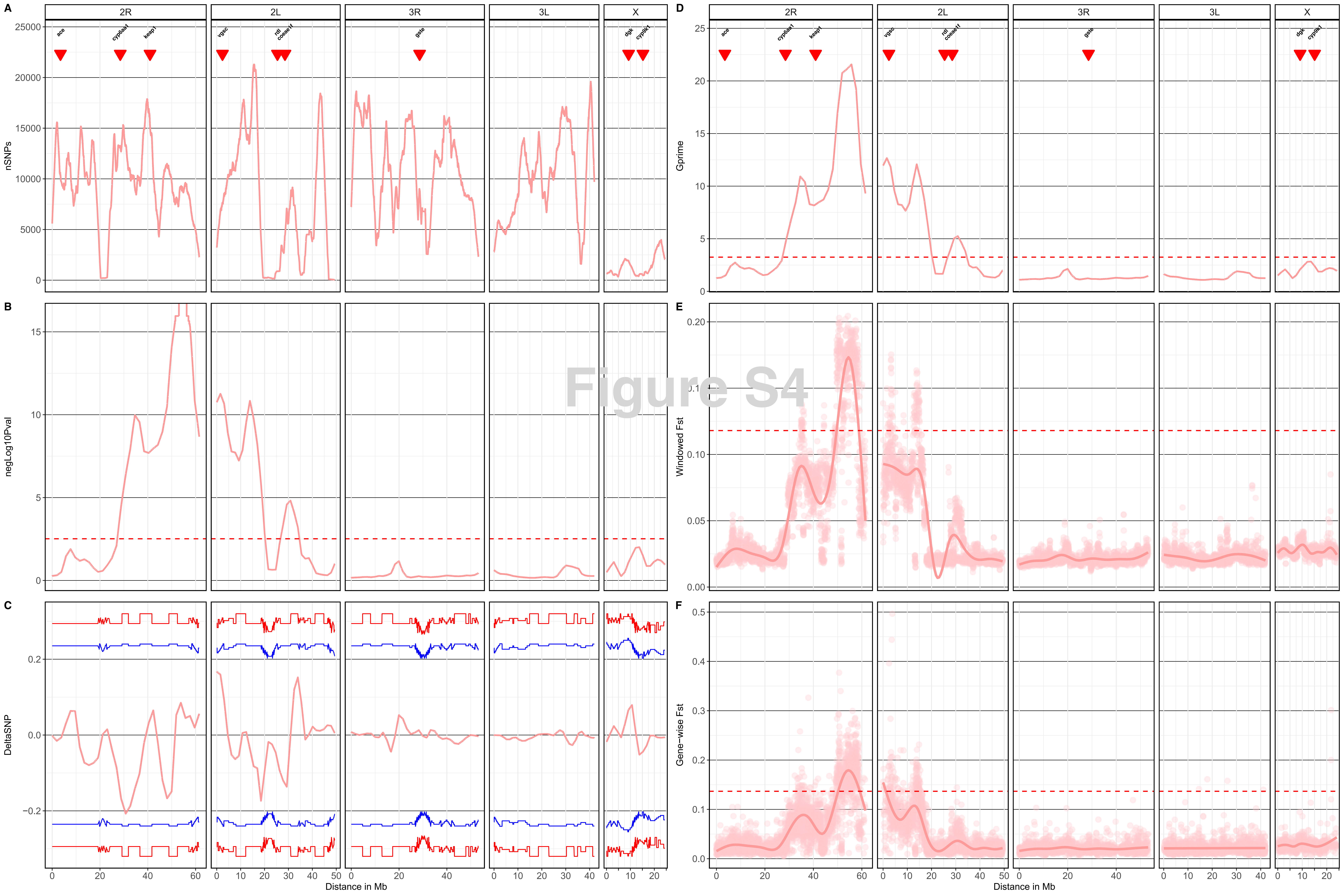
A
B
C
D
E
F
2R
2L
3R
3L
X
ace
cyp6aa1
keap1
vgsc
rdl
coeae1f
gste
dgk
cyp9k1
nSNPs
negLog10Pval
DeltaSNP
Gprime
Windowed Fst
Gene-wise Fst
Distance in Mb
Figure S4

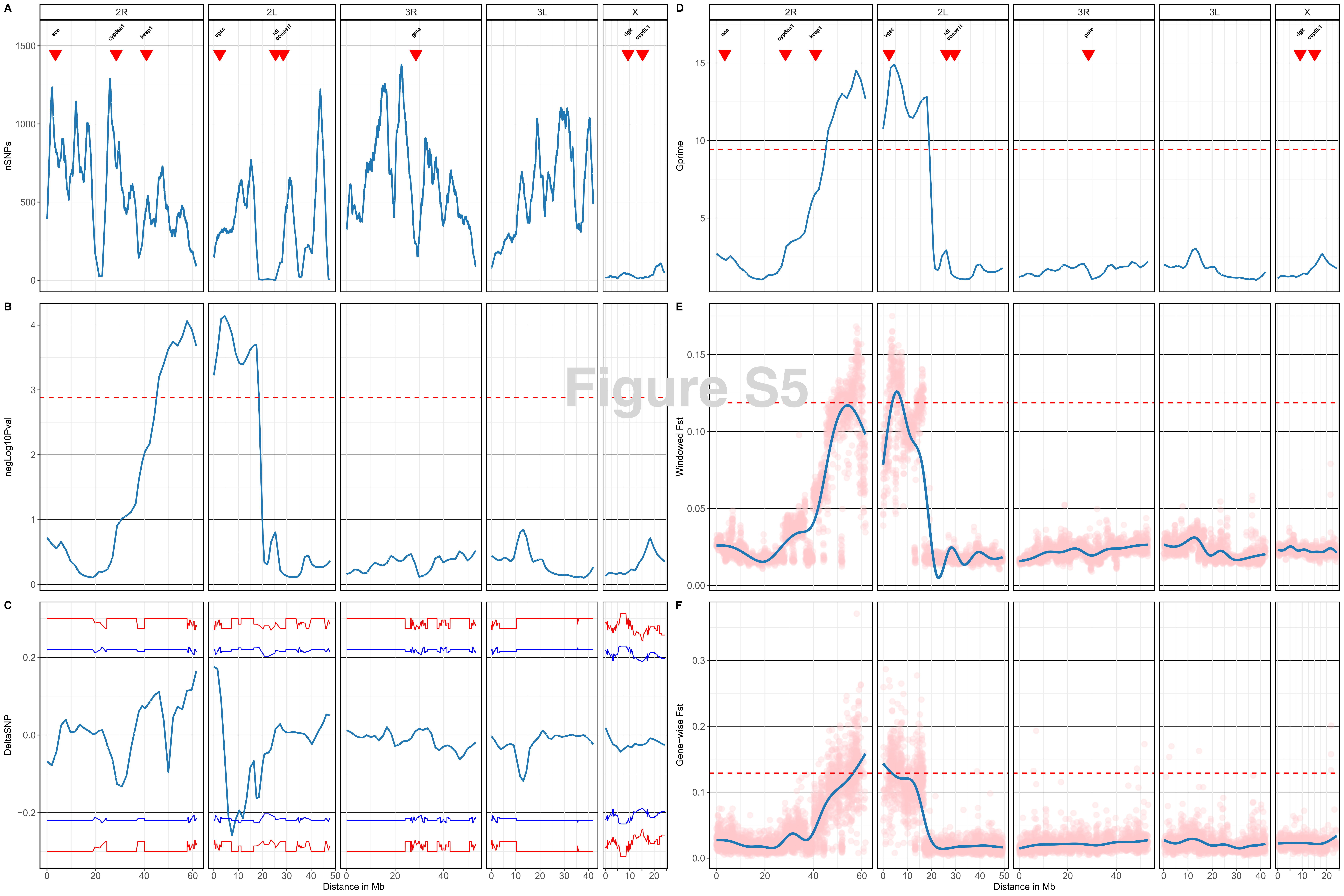

Figure S5
A
B
C
D
E
F
2R
2L
3R
3L
X
ace
cyp6aa1
keap1
vgsc
rdl
coeae1f
gste
dgk
cyp9k1
nSNPs
negLog10Pval
DeltaSNP
Gprime
Windowed Fst
Gene-wise Fst
Distance in Mb

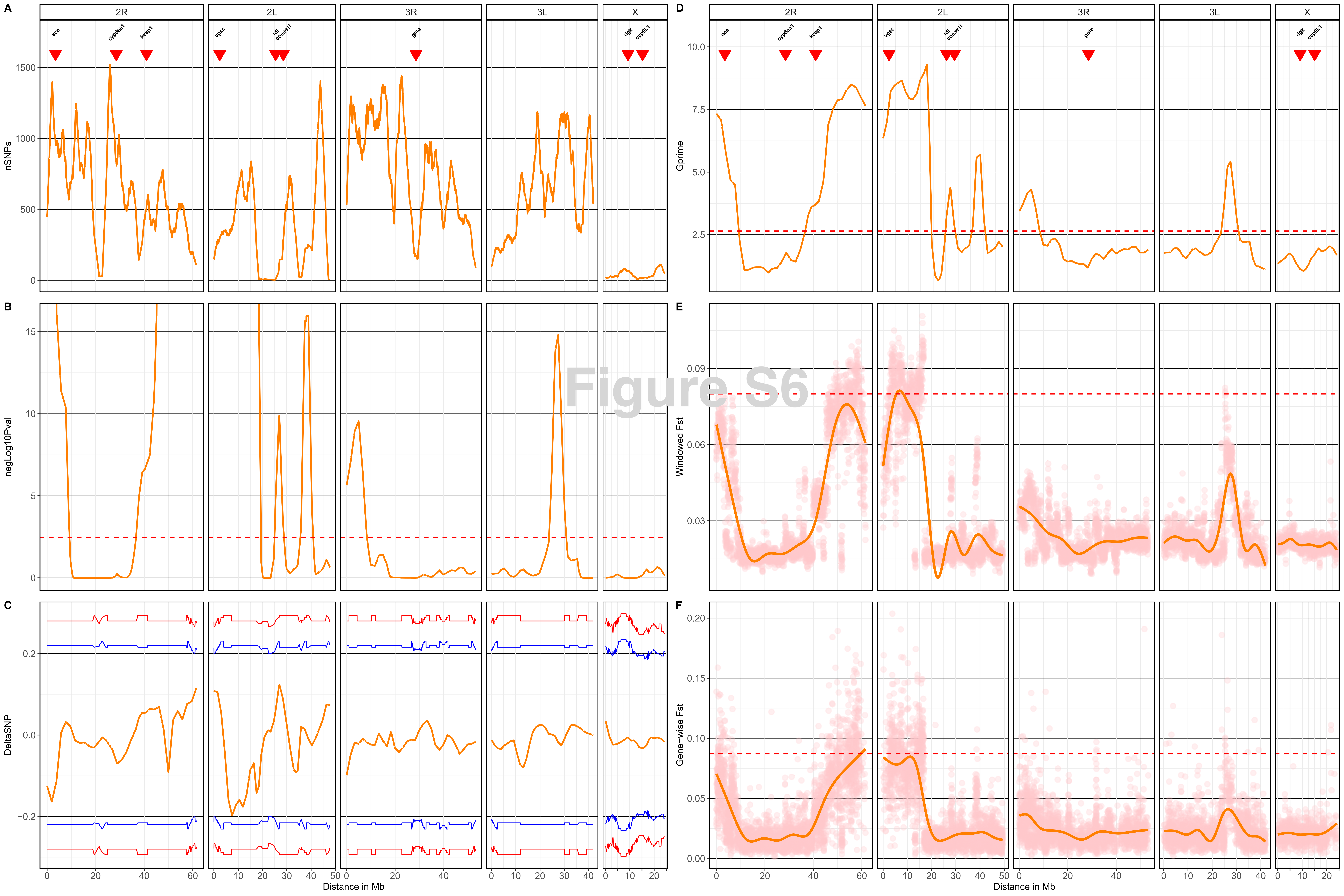

A
B
C
D
E
F
2R
2L
3R
3L
X
ace
cyp6aa1
keap1
vgsc
rdl
coeae1f
gste
dgk
cyp9k1
nSNPs
negLog10Pval
DeltaSNP
Gprime
Windowed Fst
Gene−wise Fst
Distance in Mb
Figure S6

#### vgsc Missense variant frequencies

| | AL-DDT-A | DE-DDT-A | AL-DDT-B | DE-DDT-B | AL-Perm-A-30 | AL-Perm-A-25 | DE-Perm-A-3 | AL-Perm-A-8 | AL-Perm-B-75 | AL-Perm-B-60 | DE-Perm-B-15 | DE-Perm-B-20 | Codon | AA | Sigificance |
|---|---|---|---|---|---|---|---|---|---|---|---|---|---|---|---|
| para 2422651 T>C | 0.02 | 0.06 | 0.03 | 0.13 | 0.46 | 0.5 | 0.03 | 0.17 | 0.81 | 0.7 | 0.24 | 0.46 | 3023T>C | L995S | ● (Perm-A AL30vsDE8) |
| para 2400071 G>A | 0.1 | 0.15 | 0.78 | 0.48 | 0.3 | 0.38 | 0.58 | 0.55 | 0.22 | 0.31 | 0.76 | 0.62 | 1488G>A | M490I | ● (Perm-A AL25vsDE3) ● (Perm-A AL30vsDE8) |
| para 2424759 G>T | 0.06 | 0.08 | 0.12 | 0.06 | 0.12 | 0.12 | 0.1 | 0.07 | 0.08 | 0.09 | 0.14 | 0.09 | 3556G>T | G1173C | |
| para 2399929 C>A | 0.05 | 0.12 | 0.06 | 0.08 | 0.03 | 0.11 | 0.1 | 0.09 | 0.07 | 0.02 | 0.1 | 0.08 | 1346C>A | A443D | |
| para 2424780 T>A | 0.05 | 0.05 | 0.09 | 0.06 | 0.1 | 0.08 | 0.07 | 0.04 | 0.03 | 0.02 | 0.08 | 0.06 | 3577T>A | F1180I | |
| para 2424735 G>T | 0.03 | 0.02 | 0.14 | 0.1 | 0.1 | 0.11 | 0.04 | 0.07 | 0.02 | 0.04 | 0.07 | 0.06 | 3532G>T | D1165Y | |
| para 2359643 G>T | 0.05 | 0 | 0 | 0 | 0 | 0.11 | 0 | 0.06 | 0 | 0.05 | 0.16 | 0.07 | 151G>T | D47Y | |
| para 2424785 G>T | 0.04 | 0.05 | 0.07 | 0.02 | 0.02 | 0.02 | 0.02 | 0.02 | 0 | 0 | 0.1 | 0.09 | 3582G>T | K1181N | |
| para 2399943 C>A | 0.02 | 0.06 | 0.05 | 0.06 | 0 | 0.02 | 0.02 | 0.02 | 0.02 | 0.02 | 0.02 | 0.1 | 1360C>A | Q448K | |
| para 2424754 C>A | 0.07 | 0.05 | 0.12 | 0.04 | 0.05 | 0.02 | 0.06 | 0.05 | 0.03 | 0.02 | 0.07 | 0.02 | 3551C>A | S1171Y | |
| para 2399948 A>C | 0.05 | 0.04 | 0.13 | 0.04 | 0 | 0.07 | 0.05 | 0.02 | 0.05 | 0.02 | 0.05 | 0.02 | 1365A>C | Q449H | |
| para 2417648 C>A | 0.09 | 0.07 | 0.04 | 0 | 0.11 | 0.03 | 0 | 0 | 0 | 0.12 | 0.04 | 0.03 | 2688C>A | F883L | |
| para 2424796 A>C | 0.02 | 0 | 0 | 0.02 | 0 | 0.04 | 0.07 | 0 | 0.03 | 0 | 0 | 0 | 3593A>C | H1185P | |
| para 2417689 C>A | 0 | 0.03 | 0 | 0 | 0 | 0.05 | 0.06 | 0 | 0 | 0.02 | 0 | 0 | 2729C>A | S897Y | |
| para 2417651 G>T | 0 | 0 | 0 | 0 | 0.04 | 0.06 | 0.07 | 0 | 0.03 | 0 | 0 | 0.03 | 2691G>T | K884N | |
| para 2407939 A>C | 0 | 0 | 0 | 0.04 | 0 | 0.06 | 0 | 0 | 0 | 0 | 0 | 0 | 2101A>C | N688H | |
| para 2424753 T>A | 0.02 | 0.03 | 0.02 | 0.02 | 0.02 | 0.02 | 0.02 | 0.02 | 0.05 | 0.04 | 0.02 | 0 | 3550T>A | S1171T | |
| para 2391308 A>C | 0.02 | 0.02 | 0.01 | 0 | 0.01 | 0.02 | 0 | 0 | 0.04 | 0.05 | 0.06 | 0.02 | 1302A>C | E428D | |
| para 2424391 G>T | 0.02 | 0.05 | 0 | 0 | 0 | 0.04 | 0.02 | 0.04 | 0.02 | 0.02 | 0.06 | 0.04 | 3420G>T | K1127N | |
| para 2424739 C>A | 0 | 0.03 | 0.08 | 0.02 | 0.02 | 0.04 | 0.06 | 0 | 0.02 | 0 | 0.02 | 0.04 | 3536C>A | T1166N | |
| para 2399909 A>T | 0 | 0 | 0.04 | 0.02 | 0.02 | 0.05 | 0.02 | 0 | 0 | 0.02 | 0 | 0.06 | 1326A>T | E436D | |
| para 2424384 C>A | 0 | 0.05 | 0.03 | 0 | 0 | 0.02 | 0.02 | 0.06 | 0 | 0.05 | 0.02 | 0 | 3413C>A | A1125E | |
| para 2417001 C>A | 0 | 0 | 0.03 | 0 | 0 | 0 | 0 | 0.07 | 0 | 0 | 0 | 0 | 2432C>A | T798K | |
| para 2428105 A>C | 0.04 | 0 | 0 | 0.02 | 0.02 | 0.04 | 0.02 | 0 | 0 | 0.07 | 0 | 0.05 | 4033A>C | M1332L | |
| para 2424775 G>A | 0.02 | 0 | 0 | 0 | 0.06 | 0.04 | 0 | 0 | 0 | 0.02 | 0.02 | 0.02 | 3572G>A | R1178H | |
| para 2422587 T>A | 0 | 0 | 0 | 0 | 0 | 0 | 0 | 0 | 0 | 0.06 | 0 | 0 | 2959T>A | C974S | |
| para 2407964 C>A | 0 | 0 | 0 | 0 | 0 | 0 | 0 | 0 | 0 | 0.09 | 0 | 0 | 2126C>A | A696D | |
| para 2417030 A>C | 0 | 0 | 0.03 | 0 | 0 | 0 | 0 | 0 | 0 | 0.06 | 0.02 | 0 | 2461A>C | M808L | |
| para 2416906 T>G | 0 | 0 | 0.05 | 0 | 0 | 0 | 0 | 0 | 0 | 0 | 0.02 | 0 | 2337T>G | C766W | |
| para 2381092 G>T | 0 | 0 | 0.06 | 0 | 0 | 0 | 0 | 0 | 0 | 0 | 0.02 | 0.02 | 331G>T | D105Y | |
| para 2417026 C>A | 0.02 | 0 | 0.07 | 0 | 0 | 0 | 0 | 0 | 0 | 0 | 0 | 0 | 2457C>A | H806Q | |
| para 2417011 G>T | 0.02 | 0 | 0.07 | 0 | 0 | 0 | 0 | 0 | 0 | 0 | 0 | 0 | 2442G>T | M801I | |
| para 2400038 G>T | 0.02 | 0 | 0.05 | 0 | 0.02 | 0.02 | 0 | 0 | 0.02 | 0 | 0 | 0 | 1455G>T | E479D | |
| para 2417666 G>T | 0.04 | 0 | 0 | 0 | 0 | 0 | 0 | 0 | 0.07 | 0 | 0 | 0 | 2706G>T | W889C | |
| para 2424721 C>A | 0.05 | 0.04 | 0 | 0.04 | 0.04 | 0.02 | 0.06 | 0 | 0.03 | 0 | 0 | 0 | 3518C>A | S1160Y | |
| para 2424303 G>T | 0.05 | 0.02 | 0 | 0.03 | 0 | 0 | 0.04 | 0 | 0 | 0.04 | 0 | 0.02 | 3332G>T | G1098V | |
| para 2422762 C>T | 0.06 | 0.06 | 0.03 | 0.07 | 0 | 0 | 0 | 0 | 0 | 0 | 0 | 0 | 3077C>T | S1013F | |
| para 2407925 G>T | 0 | 0.03 | 0.04 | 0.07 | 0 | 0 | 0 | 0 | 0 | 0 | 0 | 0 | 2087G>T | G683V | |
| para 2417679 C>A | 0.04 | 0.11 | 0 | 0.02 | 0.03 | 0 | 0.07 | 0.03 | 0.03 | 0.02 | 0 | 0 | 2719C>A | L894I | |
| para 2391247 T>G | 0.07 | 0 | 0.04 | 0 | 0.04 | 0 | 0.01 | 0.07 | 0 | 0 | 0.06 | 0 | 1241T>G | I408S | |
| para 2359650 G>T | 0.11 | 0 | 0 | 0 | 0 | 0 | 0 | 0.07 | 0 | 0 | 0 | 0 | 158G>T | G49V | |
| para 2431417 A>G | 0.37 | 0.26 | 0 | 0 | 0 | 0 | 0 | 0 | 0 | 0 | 0 | 0 | 6187A>G | I2050V | |
| para 2359663 G>T | 0.19 | 0 | 0.14 | 0.08 | 0.25 | 0.12 | 0 | 0.09 | 0.09 | 0.07 | 0.29 | 0.08 | 171G>T | K53N | |

Significant in:
- ● Perm-A AL25vsDE3
- ● Perm-A AL30vsDE8

Frequency: 0.8, 0.6, 0.4, 0.2, 0.0

### Figure S7
